# Tissue-aware RNA-Seq processing and normalization for heterogeneous and sparse data

**DOI:** 10.1101/081802

**Authors:** Joseph N. Paulson, Cho-Yi Chen, Camila M. Lopes-Ramos, Marieke L Kuijjer, John Platig, Abhijeet R. Sonawane, Maud Fagny, Kimberly Glass, John Quackenbush

## Abstract

Although ultrahigh-throughput RNA-Sequencing has become the dominant technology for genome-wide transcriptional profiling, the vast majority of RNA-Seq studies typically profile only tens of samples, and most analytical pipelines are optimized for these smaller studies. However, projects are generating ever-larger data sets comprising RNA-Seq data from hundreds or thousands of samples, often collected at multiple centers and from diverse tissues. These complex data sets present significant analytical challenges due to batch and tissue effects, but provide the opportunity to revisit the assumptions and methods that we use to preprocess, normalize, and filter RNA-Seq data – critical first steps for any subsequent analysis. We find analysis of large RNA-Seq data sets requires both careful quality control and that one account for sparsity due to the heterogeneity intrinsic in multi-group studies. An R package instantiating our method for large-scale RNA-Seq normalization and preprocessing, YARN, is available at bioconductor.org/packages/yarn.

**Highlights:**

- Overview of assumptions used in preprocessing and normalization
- Pipeline for preprocessing, quality control, and normalization of large heterogeneous data
- A Bioconductor package for the YARN pipeline and easy manipulation of count data
- Preprocessed GTEx data set using the YARN pipeline available as a resource

## Introduction

RNA-Seq experiments using ultrahigh-throughput sequencing-by-synthesis technologies were first performed in 2008 and have since been used for large-scale transcriptome analysis and transcript discovery in mammalian genomes (Lister et al., 2008; Mortazavi et al., 2008; Nagalakshmi et al., 2008). Although hundreds of published studies have used this technology to assay gene expression, the majority consist of a relatively small numbers of samples. There are many widely used methods for normalization and analysis of expression data from modest numbers of relatively homogeneous samples (Bolstad et al., 2003; Eisenberg and Levanon, 2013; Vandesompele et al., 2002). The workflow for RNA-Seq typically includes basic quality control on the raw reads and alignment of those reads to a particular reference database to extract sequence read counts for each feature – gene, exon, or transcript – being assayed (Conesa et al., 2016). The resulting features-by-samples matrix is then filtered and normalized and analyzed to identify features that are differentially expressed between phenotypes or conditions. Functional enrichment analysis is then performed on these features (Conesa et al., 2016).

However, there are now many large cohort studies, including the Genotype-Tissue Expression project (GTEx) and The Cancer Genome Atlas (TCGA) that have generated transcriptomic data on large populations and across multiple tissues or conditions to study patterns of gene expression (Ardlie et al., 2015; McLendon et al., 2008). The GTEx project is collecting genome-wide germline SNP data and gene expression data from an array of different tissues on a large cohort of research subjects. GTEx release version 6.0 sampled over 500 donors with phenotypic information representing 9,435 RNA-Seq assays performed on 53 conditions (51 tissues and two derived cell lines) we chose to analyze. GTEx assayed expression in 30 tissue types, which were further divided these into 53 tissue subregions (Ardlie et al., 2015).

After removing tissues with very few samples (less than 15) this led to 27 tissue types, from 49 subregions. This included 13 different brain regions and three types of skin tissue. While GTEx was very broad, the sampling is uneven across these subregions, with some sampled in nearly every donor and others sampled in only a small subset. For example, there are some tissues, such as the brain, in which many subregions were sampled with the expectation that those samples might exhibit very different patterns of expression.

Established methods for RNA-Seq analysis can be used to make direct comparisons of gene expression profiles between phenotypic groups within a tissue. However, they are not well suited for comparisons across multiple, diverse tissues that each exhibit a combination of commonly expressed and tissue-specific genes. This characteristic is a feature that confounds most normalization methods that assume the majority of expressed transcripts are common across samples. Standard normalization methods make assumptions that are valid only in fairly consistent samples, e.g. most genes are not differentially expressed, housekeeping genes are expressed at equivalent rates, or that the expression distributions vary only slightly due to technology (Bolstad et al., 2003; Eisenberg and Levanon, 2013; Vandesompele et al., 2002). In large heterogeneous data like GTEx these biological assumptions are violated when looking at completely different tissues and new methods with valid assumptions are necessary for valid comparisons.

**Figure 1:**
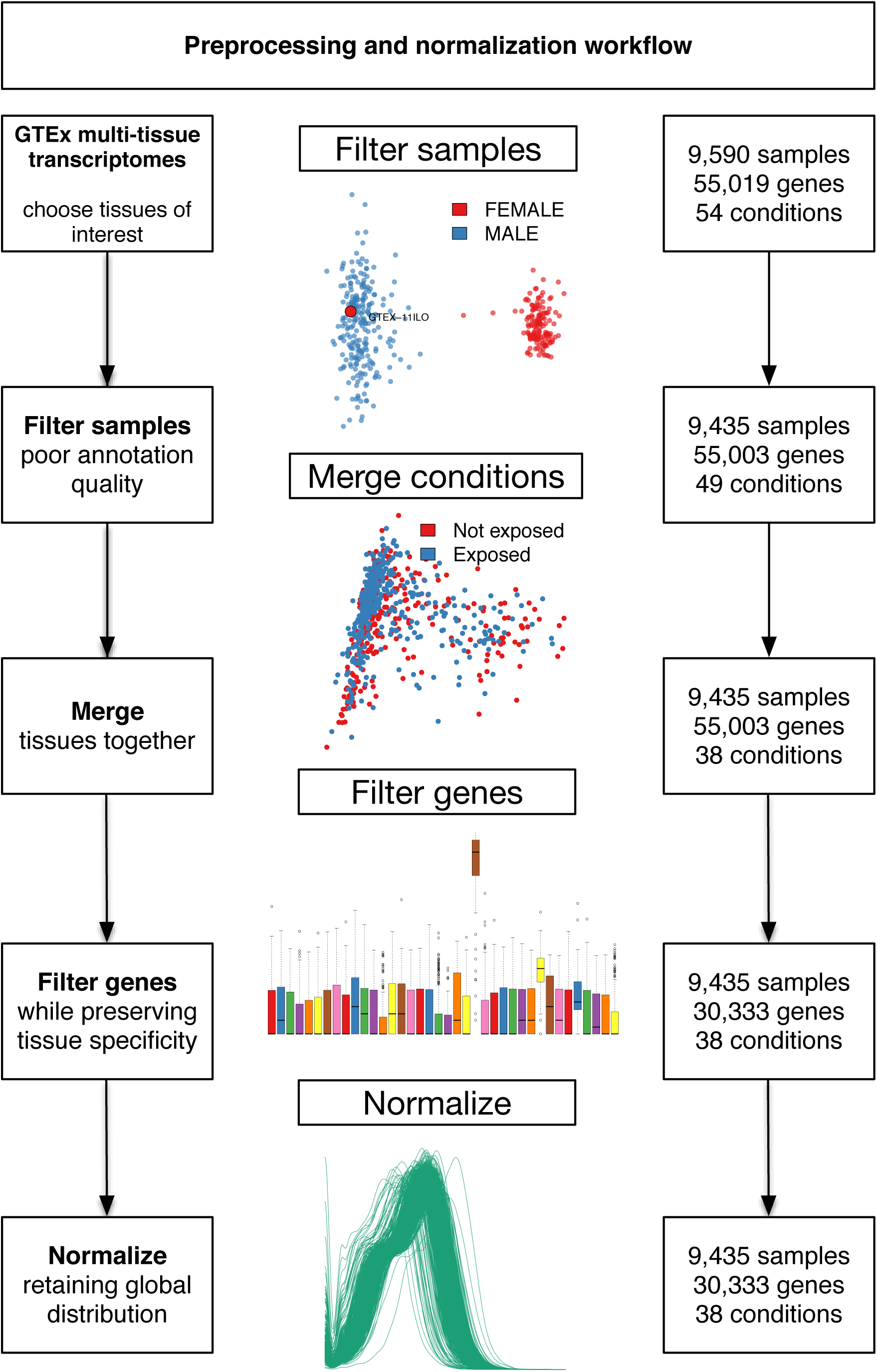
Preprocessing workflow for large, heterogeneous RNA-Seq data sets, as applied to the GTEx data. The boxes on the right show the number of samples, genes, and tissue types at each step. First, samples were filtered using PCoA with Y-chromosome genes to test for correct annotation of the sex of each sample. PCoA was used to group or separate samples derived from related tissue regions. Genes were filtered to select a normalization gene set to preserve robust, tissue-dependent expression. Finally, the data were normalized using a global count distribution method to support cross-tissue comparison while minimizing within-group variability.

Our approach, YARN, shown in Figure 1, includes filtering poorly annotated samples, merging samples representing similar information (with indistinguishable expression profiles), filtering genes in a condition specific manner, and normalizing to keep global distributions while controlling for within group-variability. An R package instantiating our method for large-scale RNA-Seq preprocessing and normalization is available at bioconductor.org/packages/yarn.

## A Robust Multi-tissue Normalization Pipeline

### Annotation Quality Assessment

The first step in any good data processing pipeline is quality assessment to assure that samples are appropriately labeled. Reliable metadata is critical for studies and a high rate of mis-assignment raises issues about the quality of the rest of the annotation provided for each sample. Some disease states and sex annotation metadata can be checked with the RNA-Seq expression values using disease biomarkers or sex chromosomal genes. Mis-annotation is a rampant issue with 46% of studies potentially having had poor annotation (Toker et al., 2016). We ourselves have needed to remove 6% of samples in an analysis of sexual dimorphism in COPD due to potential mis-annotation (Glass et al., 2014). While correct sex assignment is not a guarantee that the rest of the annotation is correct it is a good indicator that the study was thorough.

GTEx collected tissue from the autopsy of subjects with heterogeneous causes of death. The only thing we could look at was sex. We extracted count values for genes mapped to the Y chromosome in each sample and log_2_-transformed the data and used Principal Coordinate Analysis (PCoA) with Euclidean distance, a method which is equivalent to standard PCA, to cluster individuals within each tissue. While the outcome is the same for PCoA and PCA, the distance between two samples allows for an intuitive interpretation of the quality and reproducibility of a sample and any appropriate distance can be substituted. For example, correlation might not be ideal, as it cannot detect cases where there are large average shifts in expression.

PCoA clearly separates samples into two groups in every tissue using the Y chromosome genes. However, one subject, GTEX-11ILO, as female, grouped with males in each of the 12 tissue regions for which RNA-Seq data was collected (Figure S1). We excluded this sample from further analysis.

### Merging or Splitting Sample Groups

GTEx sampled 51 body sites based on morphological definitions and created two cell lines, but they did not examine whether these groups were fundamentally different in expression. To improve power for differential expression, eQTL, and gene regulatory network construction we investigated if samples from similar body regions could be grouped together due to similar expression values. We used dimension reduction approaches, specifically PCoA, to see how similar samples from subregions were to each other and to determine if samples of similar regions could be merged.

We used the GTEx annotated tissue regions, labeled SMTS, to investigate sample subregion similarity. Specifically, for each tissue region sampled in GTEx, we performed a PCoA with Euclidean distance on the log_2_-transformed raw count expression data for the 1000 most variable genes after filtering X, Y, and MT chromosomal genes (see also Figure S2). Using the 1000 most variable genes instead of all genes allowed for computational efficiency. We then analyzed whether there were global expression patterns associated with samples derived from different subregions within that tissue. If we observed a separation, we repeated the process on the larger separated subregions. The results of our tissue clustering are summarized in Table 1.

In many cases, we found distinct differences between tissue subregions, such as the various arterial or esophageal subregions. However, in other tissues, such as sun-exposed and non-exposed skin, we found no distinguishable difference (Figure 2, Table 1) with results as well. The most pronounced consolidation effect was seen for the brain tissue, which was annotated in GTEx as representing 13 independent subregions. We found that the samples from cerebellum and cerebellar hemisphere subregions were did not separate from each other in the dimension reduction, but were distinct from the other regions. Merging the cerebellum and cerebellar hemisphere subregions (brain cerebellum) we found that basal ganglia (brain cerebellum) and other subregions (brain other) clustered into distinct groups (Figure S2).

**Figure 2:**
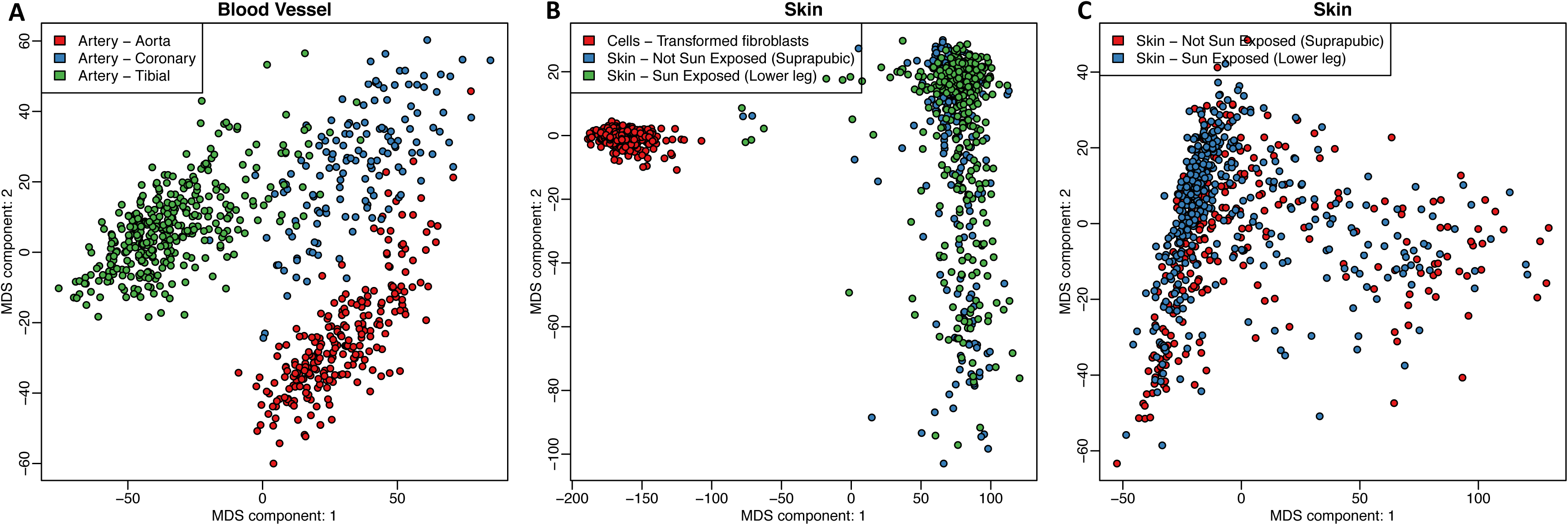
PCoA analysis allows for grouping of subregions for greater power. Scatterplots of the first and second principal coordinates from principal coordinate analysis on major tissue regions. (A) Aorta, coronary artery, and tibial artery form distinct clusters. (B) Skin samples from two regions group together but are distinct from fibroblast cell lines, a result that holds up (C) when removing the fibroblasts.

**Table 1.** Breakdown of tissues, groups we defined, abbreviations used, and sample sizes.

The PCoA clearly separated the cell lines derived from both blood and skin from their tissue of origin (Figure 2 B and C). This is consistent with previous reports that indicate that cell line generation and growth in culture media produces profound changes in gene expression (Figure 2; (Januszyk et al., 2015; Lopes-Ramos et al.)). A detailed transcriptomic and network analysis of these cell lines and their tissues of origin is provided in (Lopes-Ramos et al.).

By merging subregions together, we increased the effective sample size of several of the tissues. In doing this we were able to provide the power needed to perform eQTL analysis in several additional tissues (Fagny et al.). This increase in power is also highly beneficial for the accurate construction of gene regulatory networks, which can be analyzed in order to gain additional insights into the biological processes active in these tissues (Chen et al.; Lopes-Ramos et al.; Sonawane et al.).

### Gene Selection and Filtering for Normalization and Testing

Most standard normalization methods adjust gene expression levels using a common gene set under the assumption that the general expression distributions are roughly the same across samples. With RNA-Seq experiments, the selection of an appropriate gene set with which to carry out normalization is more challenging because, even when comparing related samples, each sample may have a slightly different subset of expressed genes. Because of this, filtering methods are essential in preprocessing RNA-Seq data to remove noisy measurements and increase power without biasing differential expression results (Bourgon et al., 2010).

In GTEx we observe many “tissue-specific” genes, or genes observed as expressed in samples collected from a single or a small number of tissues (Supplemental Methods, Figure S3). We tested two different filtering methods. First, using a “tissue-aware” manner in an unsupervised approach recommended by Anders et al. (Anders et al., 2013). Second, filtering in a “tissue-agnostic” manner to remove genes with less than one CPM in half of all samples.

This tissue-aware method filters genes with less than one count per million (CPM) in fewer than half of the number of samples of the smallest set of related samples. This approach retained 30,333 of the 55,019 mapped transcripts in the GTEx.

Filtering in a tissue-aware manner retains genes that are tissue-specific and rarely expressed genes present in at least a few samples. Retaining only highly expressed genes present in most samples, like in the tissue-agnostic approach, ignores genes that help define a tissue and are critical to tissue-specific function.

We compared the tissue-aware approach to the tissue-agnostic filtering method. Tissue-agnostic filtering removed 14,853 genes that were only expressed in a fewer than 50% of samples for which over a third are protein-coding genes and many were tissue-specific and help differentiate subregions (Supplemental Material, Table S1, Figure S4). In contrast, “tissue-aware” filtering retains the majority of the tissue-specific genes.

In Figure 3 we show examples of genes related to tissue function or health that would have been filtered using the tissue-agnostic approach. For example, *SMCP* is a protein coding gene sparse in all tissues except for the testis where it is involved in sperm motility, linked to infertility and the tumorigenicity of cancer stem-cell populations (Figure 3 E; (Hawthorne et al., 2006; Takahashi et al., 2013)). Retaining these genes for downstream analyses is crucial for understanding the relationship between gene expression and tissue-level phenotypes and understanding their impact on the complex biological system (Sonawane et al.).

**Figure 3:**
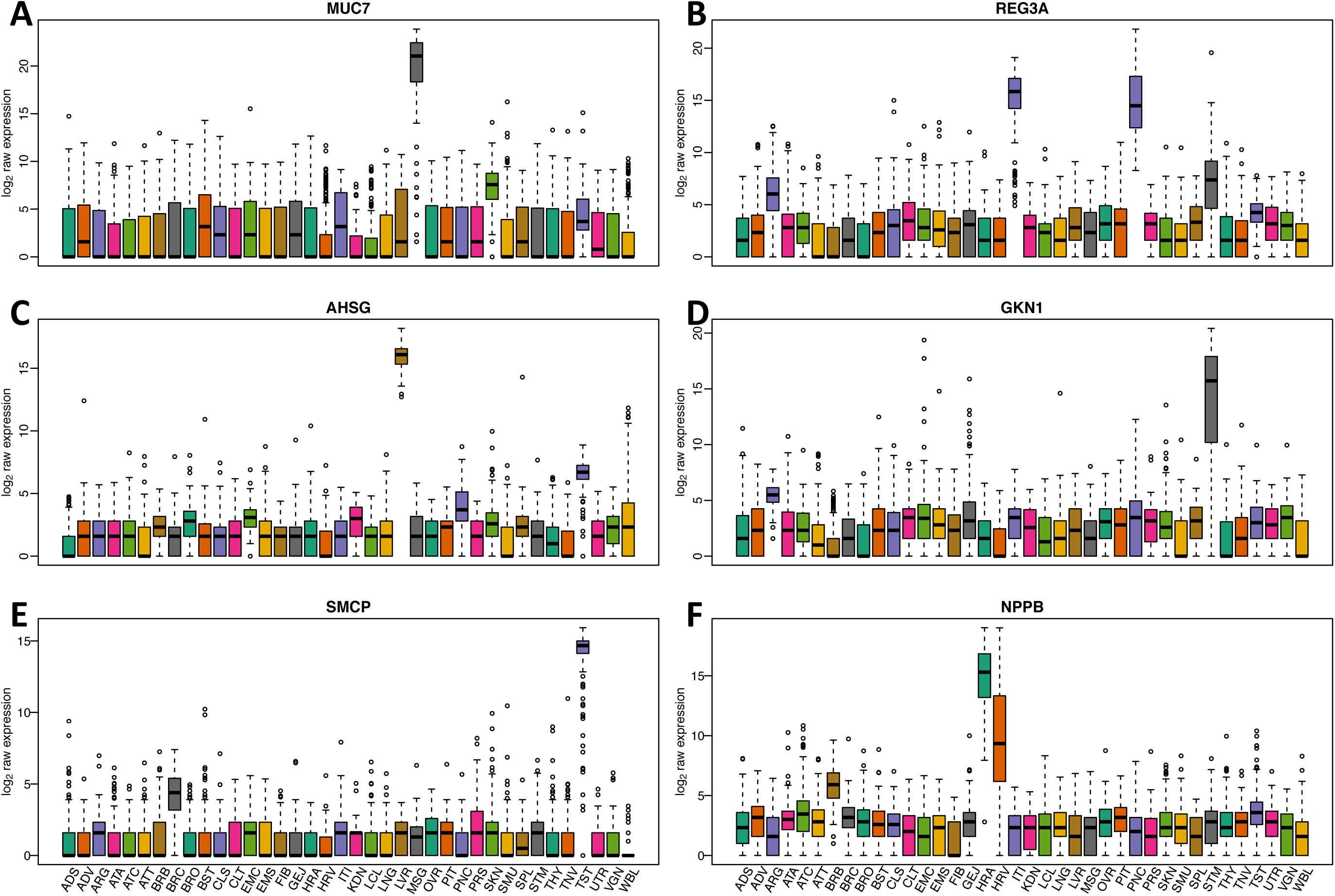
Six highly expressed tissue-specific genes that are removed upon tissue-agnostic filtering. Boxplots of continuity-corrected log2 counts for six tissue-specific genes (A-F). These genes are retained when considering tissue-specificity and not when filtering in an unsupervised manner. Colors represent different tissues. Examples include (A) *MUC7*, (B) *REG3A*, (C) *AHSG*, (D) *GKN1*, (E) *SMCP*, and (F) *NPPB*.

### Tissue-Aware Normalization

Normalization is one of the most critical steps in data preprocessing and many normalization approaches have now been developed for expression data analysis. The most widely used normalization methods for RNA-Seq were based on scaling (Anders and Huber, 2010; Bullard et al., 2010; Robinson and Oshlack, 2010), but voom introduced quantile normalization, which assumes all samples should express nearly identical sets of genes with similar distributions of expression levels. However, these assumptions break down when samples are collected from conditions under which gene expression can be expected to be different, as we see in the GTEx tissues (Figure S3, S4).

The qsmooth (Hicks et al.) normalization method is a generalization of quantile normalization that normalizes all samples together by relaxing the assumption that the statistical count distribution should be similar across all samples and only assuming it is similar in each phenotypic group (in the case of GTEx, each tissue). We applied qsmooth to the GTEx data where each phenotypic group was the merged subregions decided earlier.

Quantile normalization forces every sample’s statistical distribution to the reference’s distribution where the reference is defined as the average of all sample count quantiles. When the distributional shapes are dissimilar across tissues, the reference is not representative of any particular tissue and scaling of quantiles is dependent on the largest tissue’s distribution.

We evaluated the assumptions of quantile normalization and a relaxation by comparing the each samples’ log_2_-transformed count distribution against an all-sample reference and tissue-specific references respectively. We observed much larger root mean squared errors (RMSE) using an all-sample reference compared to tissue-specific references (Figure 4). This suggests that standard quantile normalization disproportionately weights and biases tissue-specific transcripts based on other tissues’ proportion of zeros in the distribution and tissue sample size (Supplemental Material).

**Figure 4:**
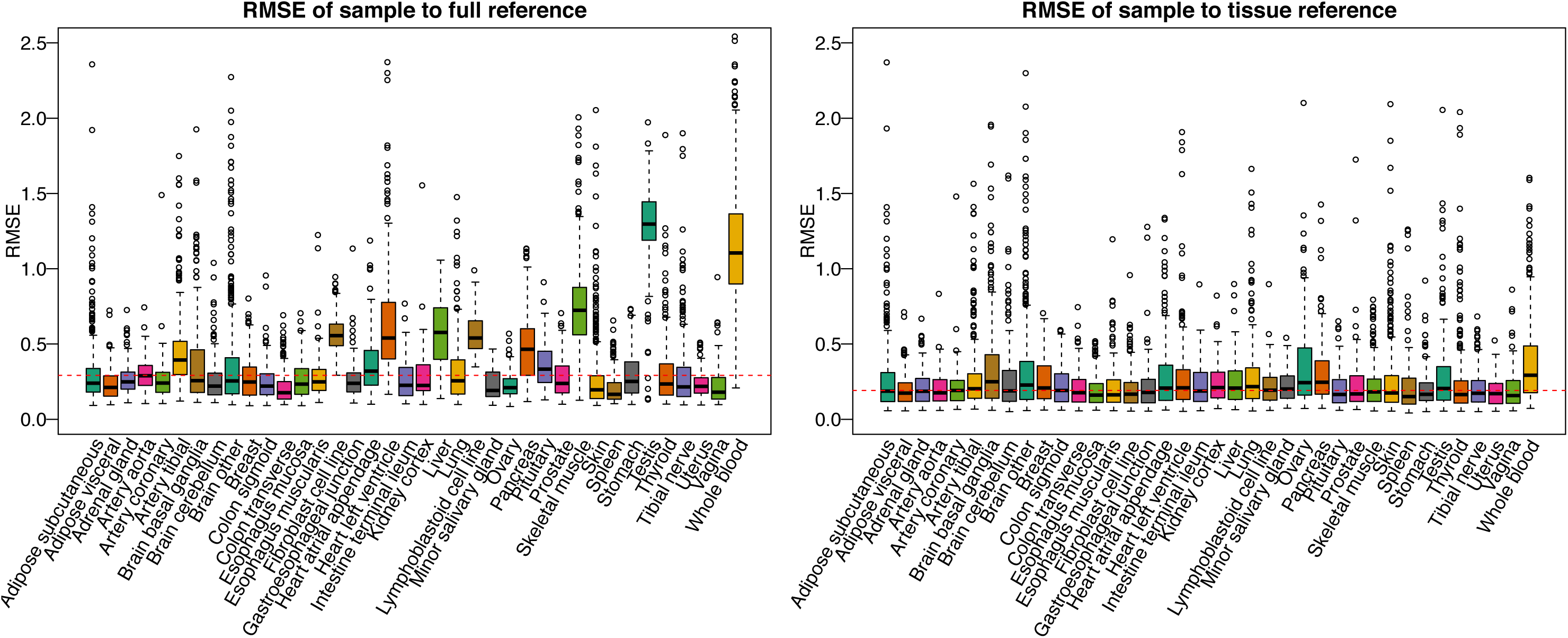
Using a tissue-defined reference lowers root mean squared error. Boxplots of the RMSE comparing the quantiles of each sample to the reference defined using (left) all tissues and samples and the (right) reference defined using samples of the same tissue.

## Conclusions

Large-scale transcriptional studies, such as GTEx, present unique opportunities to compare expression in a relatively large population and across a large number of tissues. However, as with all analyses of gene expression, it requires careful quality assessment, gene filtering, and normalization is essential if meaningful conclusions are to be drawn from the data. We developed a simple and robust pipeline, YARN, to assess annotation inaccuracies, to filter and normalize in a condition-specific manner. YARN uses PCoA (also known as classical multi-dimensional scaling) to reduce sample’s expression to two major components in search for mis-annotation on strictly phenotypic marker genes as well as on the full set of genes to see if similar conditions or tissues should be merged.

We applied YARN to the Genotype-Tissue Expression project version 6.0. YARN excluded one individual from the analysis who appears misidentified by sex and grouped many of the sampled tissue sites based on shared patterns of gene expression, increasing the effective sample size and improving the power for subsequent network analyses. In applying an optimized tissue-aware normalization process to this data, we were able to both effectively normalize commonly expressed genes and to scale tissue-specific genes appropriately. The result of our pipeline is a data set in which general expression levels are comparable between tissues, while still preserving information regarding the tissue-specific expression of genes.

YARN is freely downloadable and available under the Artistic-2.0 license on Bioconductor at bioconductor.org/packages/yarn.

## Acknowledgments

This work was supported by grants from the US National institutes of Health, including grants from the National Heart, Lung, and Blood Institute (5P01HL105339, 5R01HL111759, 5P01HL114501, K25HL133599), the National Cancer Institute (5P50CA127003, 1R35CA197449, 1U01CA190234, 5P30CA006516), and the National Institute of Allergy and Infectious Disease (5R01AI099204). Additional funding was provided through a grant from the NVIDIA foundation. This work was conducted under dbGaP approved protocol #9112 (accession phs000424.v6.p1).

## Author contributions

All authors contributed to the conception and design of the study, participated in the analysis of the data, and to writing and editing of the manuscript. JNP wrote the YARN software package. All authors read and approved the final manuscript.

## Supplemental Figure and Table Legends

**Supplemental Figure 1: PCoA analysis of multiple tissues on Y-chromosomal genes can highlight poor gender annotation, related to Figure 1 and mis-annotation section.** Scatterplots of the first and second principal components from principal component analysis on all major tissue regions. We plotted 13 tissue regions from the GTEx consortium, coloring the annotated sex of each sample. Enlarged is sample GTEX-11ILO that clusters with male samples in every tissue despite a female annotation.

**Supplemental Figure 2: PCoA analysis of multiple tissue groups, related to Figures 1, 2 and merging conditions section.** Scatterplots of the first and second principal components from principal component analysis on all major tissue groups colored by sampled region The grouping in these plots led us to either merge regions into a single group or to keep them separate. The results are summarized in Table 1.

**Supplemental Figure 3: Animated density plots of log-transformed counts when including more tissues, related to Figures 1.** GIF animation of density plots when including 10 largest sample size tissues. As more samples are included we observe more tissue-specific genes as demonstrated by the increase in a spike-mass at zero with each tissue.

**Supplemental Figure 4: Heatmap of the 15 most variable genes in the GTEx heart samples post filtering, related to Figures 1 and 3.** Heatmap of the 15 most variable genes in the GTEx heart samples. Left, top 15 genes were chosen in an unsupervised manner using the normalized gene expression after a stringent filtering in a tissue-agnostic manner. Right, the 15 most variable genes genes were chosen in an unsupervised manner using the normalized gene expression after tissue-specific filtering.

**Supplemental Figure 5: Heatmap of the 15 most variable genes in the GTEx heart samples post filtering, related to Figures 1 and 3.** Density plots of gene count distributions. left to right: log_2_ raw expression distribution of samples pre normalization; count distribution for each sample normalized in a tissue-aware manner. Colors represent different tissues.

**Supplemental Table 1: Breakdown of gene types remaining in each data set after different filtering approaches.** Filtering in a tissue-specific manner, 30,333 genes of which 60% (18,328) are classified as protein coding genes and 11% (3,220) are psuedogenes. This is in contrast to filtering by half of the data set that retains only 15,480 genes for which 84% (12,994) are protein coding, and 4% (659) are pseudogenes. However, from the full data set only 5% (1,223) of the genes filtered (24,686) were protein coding and 42% (10,446) were pseudogenes. By filtering by half of the data set, 5.4 times as many protein coding genes (6,557) were filtered and only 1.2 times (13,007) as many psuedogenes were removed.

